# ORAL HEALTH BEHAVIOUR PERCEPTION SCALE APPLIED AMONG A SAMPLE OF PORTUGUESE ADOLESCENTS

**DOI:** 10.1101/750240

**Authors:** Nélio Jorge Veiga, Odete Amaral, Maria José Correia, Inês Coelho

## Abstract

**Introduction:** The application of a scale can be particularly useful for the epidemiological studies comparing different populations and for analysis of the influence of distinct aspects of oral health on the development of certain health conditions. The aim of this study consists in the creation of a scale to classify the level of perception of the oral health behaviors applicable to the sample of Portuguese adolescents.

**Materials and methods:** An observational cross-sectional study was designed with a total of 649 adolescents between the ages of 12 and 18 years old from five public schools in the Viseu and Guarda districts, in Portugal. Data was collected by the application of a self-administered questionnaire and, after analysis of data collection, the newly Universidade Católica Portuguesa (UCP) oral health perception scale was created.

**Results:** Analyzing the sample included in the present study, we verified, by the UCP oral health perception scale created, that 67.9% of the sample presented a poor perception of their oral health behaviors, 23.9% intermediate/sufficient and only 8.2% presented what is considered in the scale as having a good classification in terms of oral health behaviors perception respecting the assumptions defined for the elaboration of the present scale.

**Conclusions:** For this purpose, through the scale to classify the level of oral health behaviors applicable to the sample of portuguese adolescents, it is possible to compare the data of several samples and understand what are the most frequent oral or eating habits among adolescents.

## Introduction

In the dental literature there are few publications describing the combination of different variables to create a specific oral health scale, in particular, to assess the perception of oral health behaviours among adolescentes. The idea of representing the perception of oral health in a single numerical value is of particular interest and gives the possibility to researchers to understand in a more objective and quantitative manner epidemiological variables. (1) Also, the application of a scale can be particularly useful for the epidemiological studies comparing different populations and for analysis of the influence of distinct degrees of oral health on the development of certain health conditions. (1)

Prevention of oral health in the school context should be driven by preventive programs in society, which should aim at oral health education and literacy, as well as providing information for the development of healthy oral hygiene habits. (2) Oral health education allows to predict the behaviors and habits of children and adolescentes, from which the existence or absence of oral diseases is diagnosed and their treatment defined. Another measure of the integration of oral health in schools, in the national context, is the reformulation of school curricula together with school health teams. This integration makes it possible to define health education strategies that stimulate the capacity of the school and family in the adolescent’s oral hygiene and the promotion of healthy eating. (3,4)

In order to have an adequate intervention and a clear definition of the strategies, the assessment of oral health indicators and the development of appropriate scales are essential to evaluate the epidemiological reality from a clinical and behavioral point of view. (1)

The aim of this study consists in the creation of a scale to classify the level of perception of the oral health behaviors applicable to a sample of Portuguese adolescents.

## Materials and methods

The present study is an observational cross-sectional study and obtained approval by the Health Sciences Institute of the Universidade Católica Portuguesa and the formal authorization of the participating schools. The informed and explicit consent of the adolescents participating in the study and their legal guardians was also received. A total of 649 adolescents between the ages of 12 and 18 years old from five public schools in the Viseu and Guarda districts (Aguiar da Beira, Mundão, Abraveses, Mangualde and Satão) during the year 2017 participated in this study. All schools participated in the community oral health program “My Best Smile” developed by the Institute of Health Sciences of the Universidade Católica Portuguesa.

Data was collected by the application of a self-administered questionnaire and, after analysis of data collection, the newly UCP oral health perception scale was created.

Statistical analysis was performed using SPSS-IBM software version 24.0. In the descriptive statistical analysis, absolute and descriptive frequencies were used for variables with nominal measurement level, mean as a measure of central tendency and standard deviation as a measure of dispersion for interval variables.

## Results

The process of constituting scales was properly structured for the perception of adolescents’ oral health behaviors. Thus, the following questions / variables were selected for inclusion in the scale, and data was collected using a self-administered questionnaire. All variables were assigned a numeric index as follows.

### How do you describe your oral health?

To obtain the index of the description of their own oral health we used the following items “Very good”; “Good”; “Reasonable”; ‘Weak”; “Very weak” with a score, revealing that the lower the index, the worse the adolescent’s oral health (table 1):

**Table 1-.** Scores attributed to the variable of question 1.

| Variable | Score |
| --- | --- |
| **How do you describe your oral health?** |  |
| Very good | 4 |
| Good | 3 |
| Reasonable | 2 |
| Weak | 1 |
| Very weak | 0 |

### Do you consider yourself informed about oral hygiene / oral health?

To obtain the oral hygiene / oral health information level index we used the following items “Very”; “Reasonable”; “Little”; “Nothing” with the attribution of a score, revealing that the lower the index, the worse the adolescent’s oral health knowledge and information level (table 2):

**Table 2-.** Scores attributed to the variable of question 2.

| **Variable** | **Score** |
| --- | --- |
| **Do you consider yourself informed about oral hygiene / oral health?** |  |
| Very | 3 |
| Reasonable | 2 |
| Little | 1 |
| Nothing | 0 |

### Brushing

Based on the variables of the questionnaire “Do you brush your teeth every day?” and “How many times do you brush your teeth per day?”, a numerical index of brushing was determined based on the following empirical criteria (table 3):

**Table 3-.**
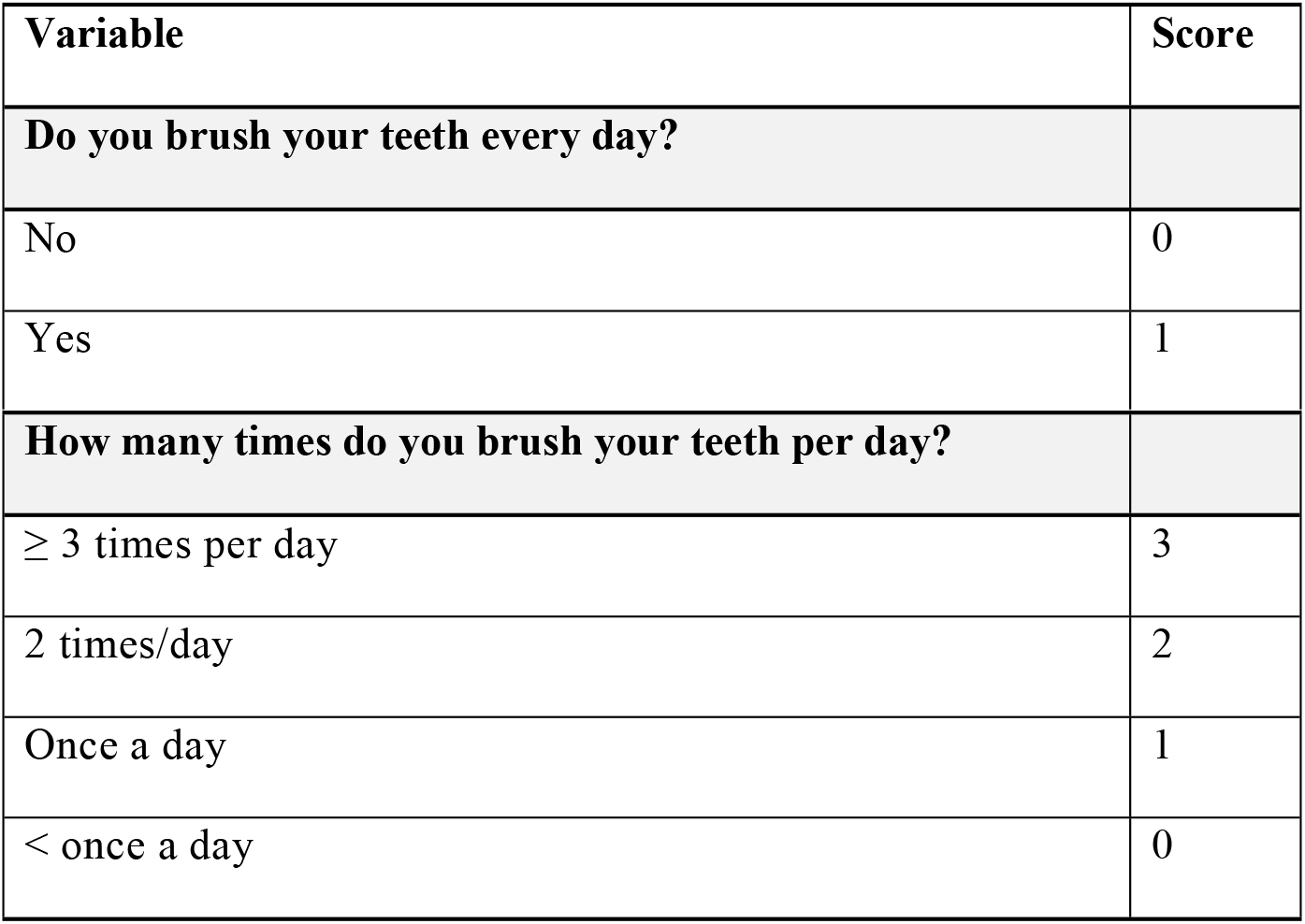
Scores attributed to the variable of questions 3 and 4.

### Average oral hygiene time

Based on the variables of the questionnaire “Each time you perform your oral hygiene on average, how long does it take?”, a numerical index of minutes spent on oral hygiene was determined based on the following empirical criteria (table 4):

**Table 4-.** Scores attributed to the variable of question 5.

| Variable | Score |
| --- | --- |
| **Average oral hygiene time.** |  |
| $\geq 3$ minutes | 3 |
| 2 minutes | 2 |
| 1 minute | 1 |
| Less than a minute | 0 |

### Achievement of oral hygiene

Based on the variables of the questionnaire “How do you perform your oral hygiene?”, A numerical index of how oral hygiene was performed based on the following empirical criteria was determined (table 5):

**Table 5-.** Scores attributed to the variable of question 6.

| Variable | Score |
| --- | --- |
| **How do you perform your oral hygiene?** |  |
| Brush gums, teeth and tongue | 3 |
| Brush teeth and tongue | 2 |
| Brush teeth and gums | 1 |
| Only brush teeth | 0 |

### Use of fluoride toothpaste

Based on the variables of the questionnaire “Do you use a fluoride toothpaste?”, a numerical index was determined taking into account the use of the toothpaste referred to in the questionnaire and based on the following empirical criteria (table 6):

**Table 6-.**
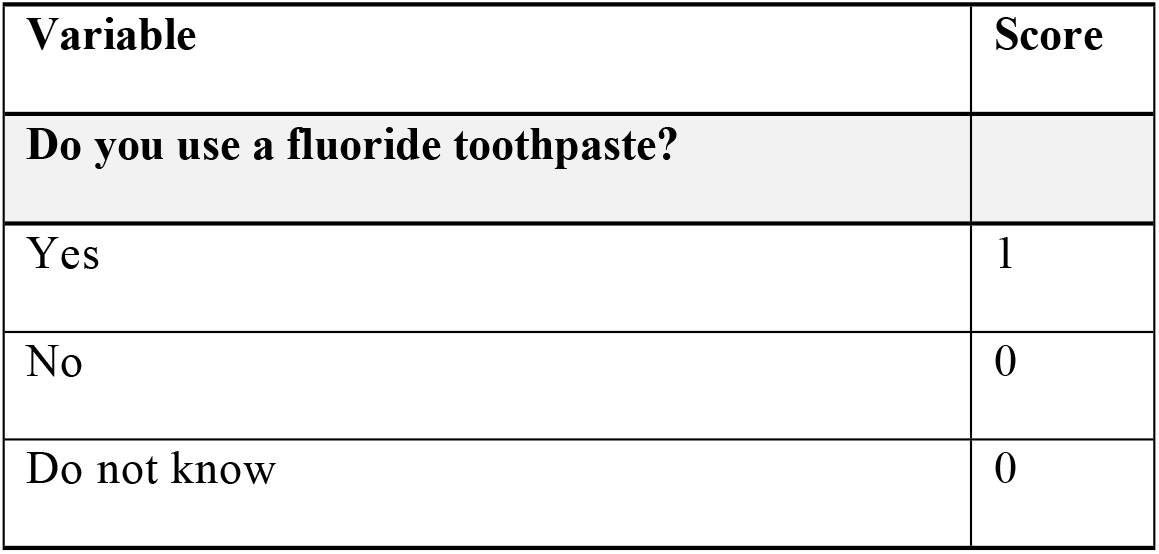
Scores attributed to the variable of question 7.

| Variable | Score |
| --- | --- |
| **Do you use a fluoride toothpaste?** |  |
| Yes | 1 |
| No | 0 |
| Do not know | 0 |

### Correct Brushing Awareness

Based on the variables of the questionnaire “With regard to brushing, do you know how to brush your teeth correctly?”, a numerical index was determined for the awareness of correct brushing based on the following empirical criteria (table 7):

**Table 7-.** Scores attributed to the variable of question 8.

| Variable | Score |
| --- | --- |
| **Do you know how to brush your teeth correctly?** |  |
| Yes | 2 |
| Reasonably | 1 |
| No | 0 |

### Flossing

Based on the variables of the questionnaire “Do you usually use dental floss?”, a numerical index regarding the use of dental floss by the adolescent was determined based on the following empirical criteria (table 8):

**Table 8-.** Scores attributed to the variable of question 9.

| Variable | Score |
| --- | --- |
| **Do you usually use dental floss?** |  |
| Yes, on a daily basis | 2 |
| Yes, sometimes | 1 |
| No | 0 |
| I do not know what is flossing | 0 |

### Dental appointment

Based on the variables of the questionnaire “Have you ever had a dental appointment?”, a numerical index was determined for the adolescent consultation with the dentist based on the following empirical criteria (table 9):

**Table 9-.** Scores attributed to the variable of question 10.

| Variable | Score |
| --- | --- |
| **Have you ever had a dental appointment?** |  |
| Yes | 1 |
| No | 0 |

### Dental appointment in the last 12 months

Based on the variables of the questionnaire “In the last 12 months have you had a dental appointment?”, a numerical index was determined for the adolescent consultation with the dentist based on the following empirical criteria (table 10):

**Table 10-.** Scores attributed to the variable of question 11.

| Variable | Score |
| --- | --- |
| **In the last 12 months have you had a dental appointment?** |  |
| Yes | 1 |
| No | 0 |
| Do not know/do not remember | 0 |

### Toothache

Based on the variables of the questionnaire “In the last 12 months, did you have a toothache?”, a numerical index was determined for the adolescent consultation with the dentist based on the following empirical criteria (table 11):

**Table 11-.**
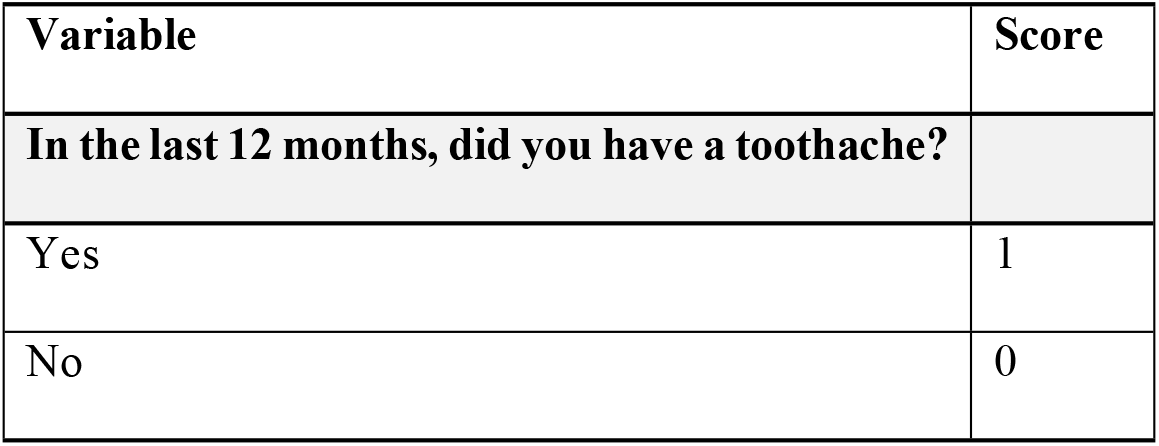
Scores attributed to the variable of question 12.

### Gum bleeding and/or pain during brushing

Based on the variables from the questionnaire “In the last 12 months did your gums bleed or hurt during tooth brushing?”, a numerical index for gingival bleeding during brushing was determined by the adolescent based on the following empirical criteria. they introduce themselves (table 12):

**Table 12-.**
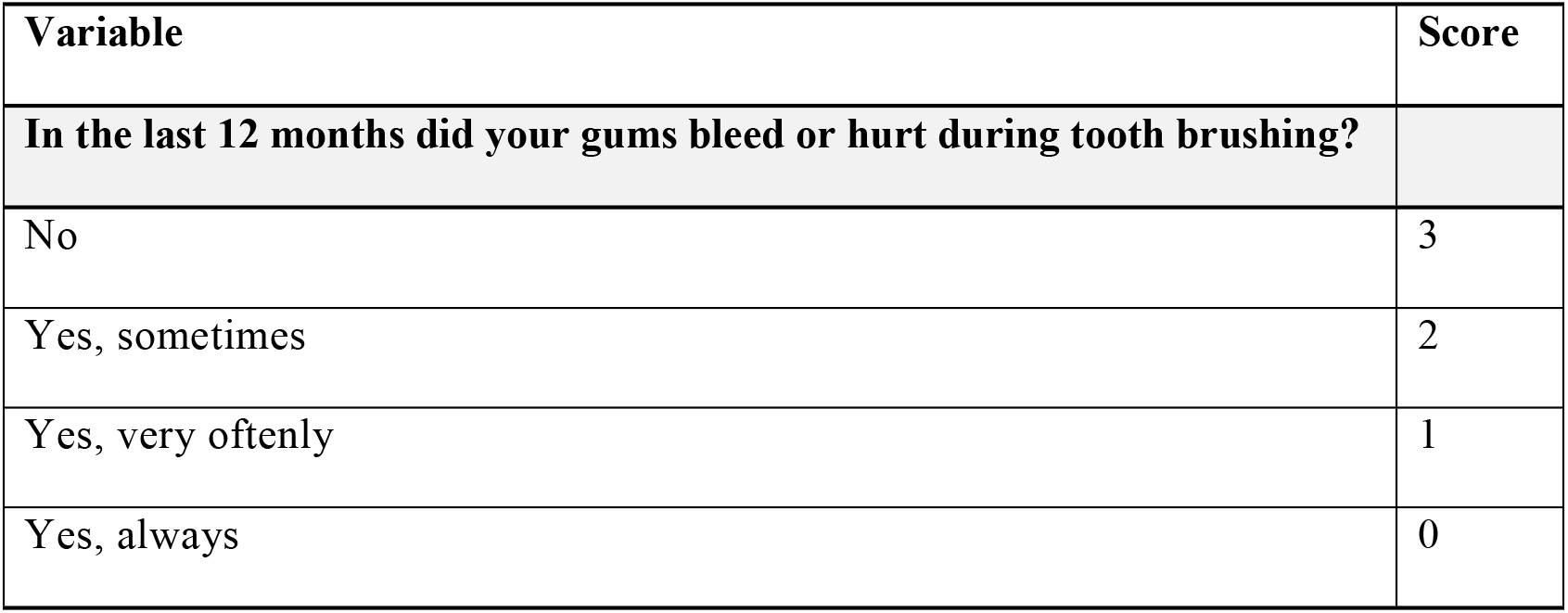
Scores attributed to the variable of question 13.

### Gum inflammation

Based on the variables of the questionnaire “In the last 12 months have you ever had inflamed (reddish) gums?”, a numerical index was determined for the presence of gingival inflammation by the adolescent based on the following empirical criteria (table 13):

**Table 13-.**
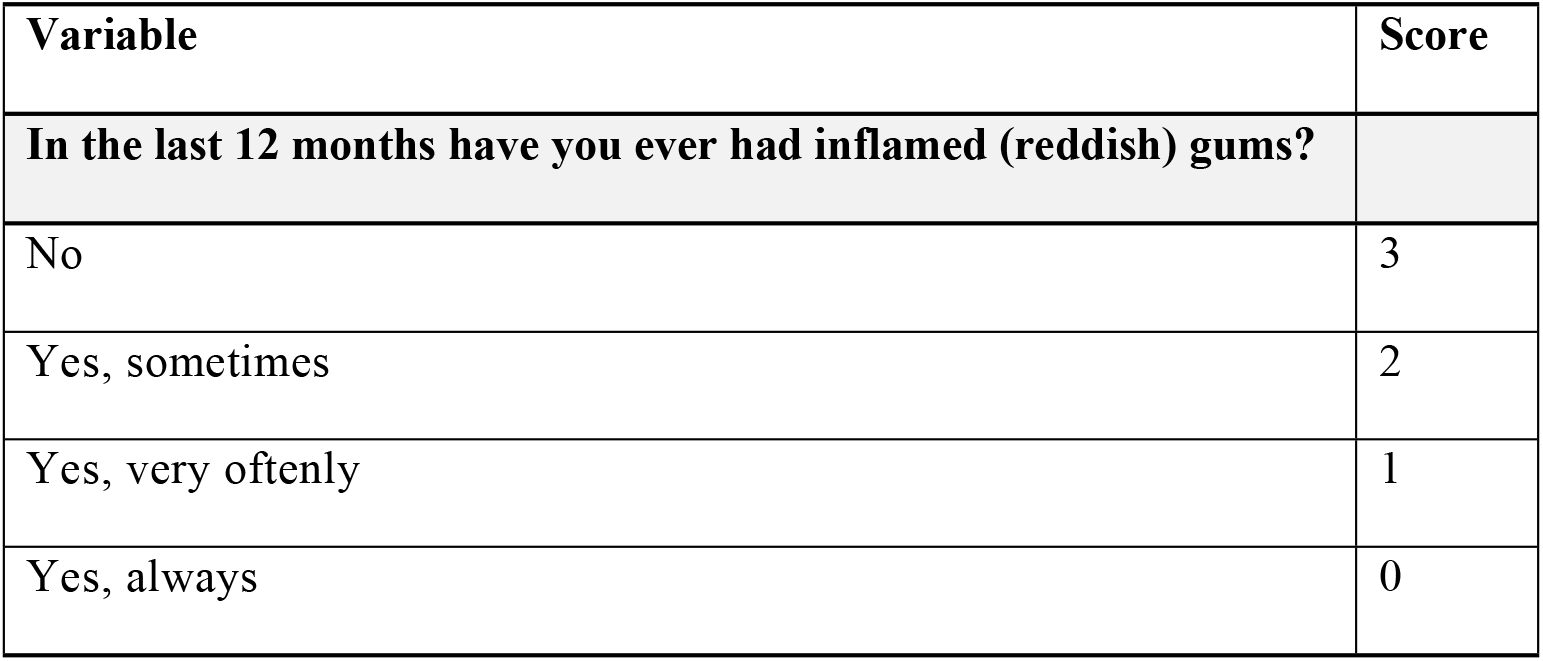
Scores attributed to the variable of question 14.

### Consumption of sugary foods

Based on the variables of the questionnaire “Do you eat sugary foods?”, a numerical index was determined for the consumption of sugary foods by adolescents based on the following empirical criteria (table 14):

**Table 14-.**
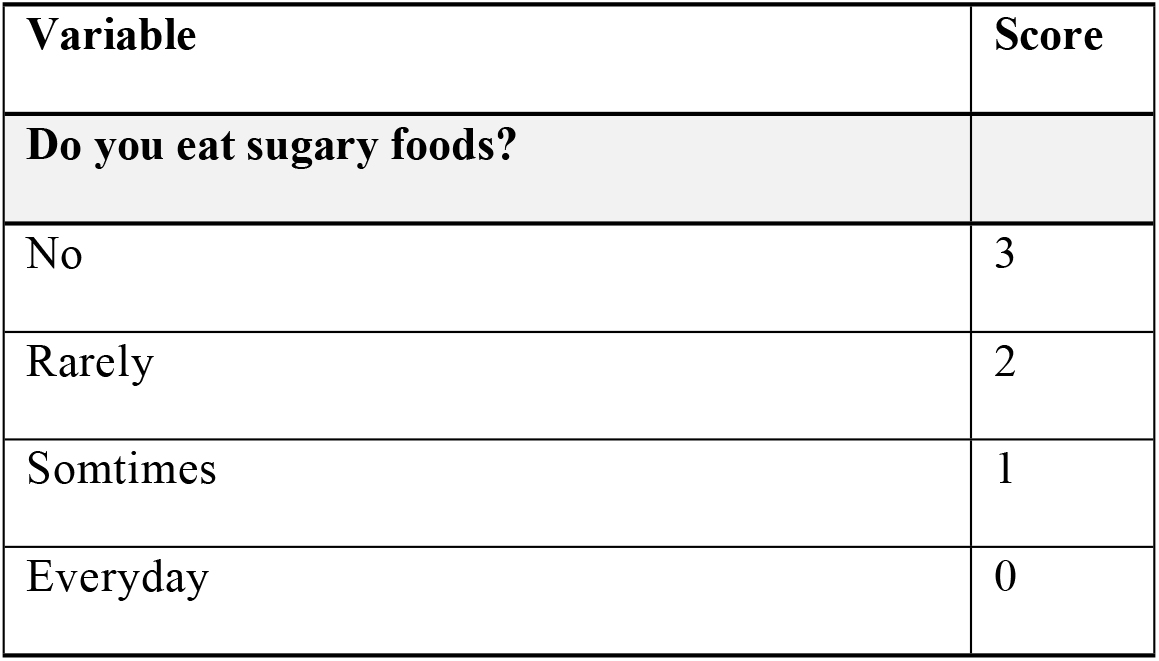
Scores attributed to the variable of question 15.

### Consumption of sugary foods between and / or after meals

Based on the variables of the questionnaire “When do you usually eat sugary foods?”, a numerical index was determined for the period of consumption of sugary foods by adolescents based on the following empirical criteria (table 15):

**Table 15-.**
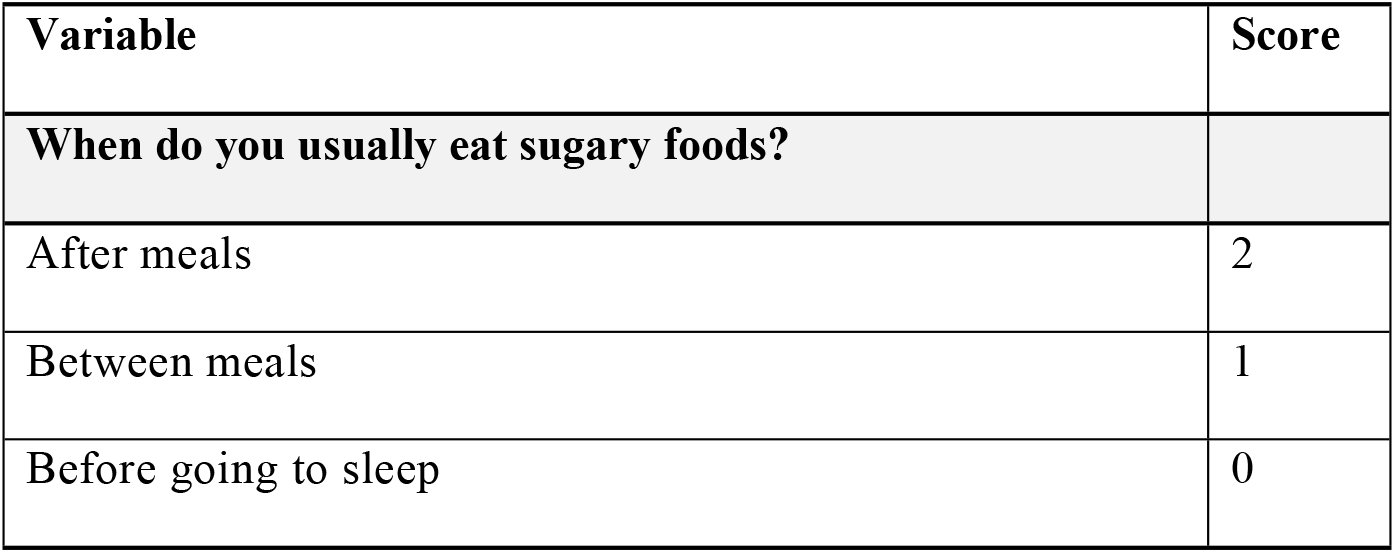
Scores attributed to the variable of question 16.

| Variable | Score |
| --- | --- |
| **When do you usually eat sugary foods?** |  |
| After meals | 2 |
| Between meals | 1 |
| Before going to sleep | 0 |

The sum of the partial indexes of the variables result in the overall oral hygiene value, which has a minimum value of 14 and a maximum of 34.

According to the response tendency, the lower this index, the less appropriate will be the oral health behaviors and the oral hygiene level of the adolescent. Based on the formula Average (M) ± 0.25 Standard Deviation (SD), cutoff groups were grouped which will allow the following classification regarding oral health behaviors in adolescents [5]:

- Insufficient - (≤ M-0.25 SD);
- Intermediate / Sufficient - (> M-0.25 SD <M + 0.25 SD);
- Good - (≥ M + 0.25 SD);

According to the data skewness and kurtosis if the groups are homogeneous, the most correct one will be used (Average±0.25 Standard Deviation), however, if they are not homogeneous groups, we can use the following formula (Median±0.25, interquartile range). From the calculations performed, we found that: Average=24.55; Standard Deviation (SD)=3.01. Thus, we obtained the following classification to determine the level of perception of oral health behaviors of a adolescent who answered the questionnaire:

- Insufficient - (≤ 23.8);
- Intermediate / sufficient - (> 23.8 and < 25.3);
- Good - (≥ 25.3).

According to the percentiles obtained:

- Percentile 25 - 23 points;
- Percentile 50 - 24.5 points;
- Percentile 75 - 27.0 points.

By the percentiles, we were able to establish the following final cutoff points for classifying the level of perception of oral health behaviors:

- Insufficient - (≤ 23.0);
- Intermediate / sufficient - (23.0 - 27.0);
- Good - (≥ 27.0).

Analyzing the sample included in the present study, we verified after applying the scale created that:

- 67.9% of the sample present insufficient perception of adequate oral health behaviors;
- 23.9% intermediate / sufficient;
- and only 8.2% present what is considered in the scale as having a good classification in terms of perception of oral health behaviors respecting the assumptions defined for the elaboration of this scale.

## Discussion

In the process of constructing the scale by the self-administered questionnaire, epidemiological data was used in order to operationalize its variables, based on the following empirical criteria:

### 1 The moderate prevalence of oral diseases, including dental caries, in the adolescent population

Dental caries in the adolescent population remains the most common chronic pathology in adolescents and is more common among children and adolescents aged 5-17 years due to poor oral hygiene, which will result in the onset of dental pathologies during childhood, youth and adulthood; at the psychological level; overall health and family quality of life. (4)

The study developed by the General Directory of Health, of Portugal, in 2008, on the prevalence of dental caries, the study demonstrated that 51% of Portuguese children with 6 years of age did not have dental caries, registering a decayed, missing and filled permanent teeth index (DMFT index) at 6, 12 and 15. years of 0.07, 1.48 and 3.04 respectively. (5)

On the other hand, the study developed by Reis *et al.* showed that the prevalence of dental caries among Portuguese 6-year-old children was 49%, rising to 72% at 15 years of age, so it is necessary to invest in the improvement oral health promotion programs for children and adolescents to prevent oral diseases. Theses community oral health interventions need to be correctly adapted to the age stage of the children and adolescents. (6)

### 2 Moderate brushing frequency at least 2 times / day

According to the study developed by the General Directory of Health, of Portugal, approximately 50% of 6-year-old children and 67-69% of 12 and 15-year-old adolescents use a toothbrush at least twice a day with a fluoride toothpaste. (6) This procedure is essential in oral hygiene as it allows removing dental plaque and food debris. To be most effective, brushing should be supplemented with the use of a fluoride toothpaste and performed at least twice a day. (7,8)

### 3 Very frequent and poor daily use of dental floss

Despite its relevance, the use of dental floss is not frequent among the young people, so its implementation and use is essential to prevent the development of dental caries on interproximal surfaces of teeth and periodontal diseases. (9,10)

### 4 Moderate appeal of the adolescent attending a dental appointment

Dental appointments should be regular or at least once every six months, which facilitates the diagnosis of oral conditions, information on the most appropriate treatments and preventive measures, such as the topical application of fluorine sealants. and fluoride. (11)

After applying the scale created, it was possible to verify that 67.9% of the sample presents a insufficient perception of oral health behaviors; 23.9% intermediate / sufficient and only 8.2% present what is considered on the scale as having a good rating in terms of the perception of oral health behaviors respecting the assumptions defined for the elaboration of this scale.

These results allow us to conclude that at school and family level, some efforts should be made, providing support and supervision at the time of oral hygiene, because only then oral hygiene will be performed frequently and properly. It was also observed that among the adolescents in this sample, despite brushing their teeth daily, they refer not using dental floss, which is essential to complement the brushing.

As recommendations for parents, caregivers and teachers, it is advisable to accompany and supervise younger children and adolescents in the prevention of oral health, particularly at the time of brushing, which should be daily, at dentist appointments of at least twice a year and provision of information for the development of healthy oral hygiene habits.

## Conclusions

For this purpose, through the scale to classify the level of perception of oral health behaviors applicable to the sample of Portuguese adolescents, it is possible to compare the data of several adolescents and understand what are the most frequent oral or eating habits among adolescents. In sum, the use of the scale is relevant, as it can be done at national, regional or local level, which provides an overview of oral and eating health habits, facilitating the area of action in the strengths and weaknesses that need to be improved, namely, oral hygiene, number of dental appointments and sugary food intake.

## Acknowledgments

The authors are deeply indebted to the teachers and students of the School Groups for the participation and important contribution to this study.

